# Flywheel Genomics: Simultaneous trait discovery and genetic gain in plant breeding

**DOI:** 10.64898/2026.08.03.742257

**Authors:** Brian Rice, Ebenezer Ogoe, Jean Rigaud Charles, Eduardo Melgar, Sandeep R. Marla, Terry Felderhoff, Allan Fritz, Geoff Morris, Gael Pressoir

**Author notes:** Corresponding authors: Brian Rice, Gael Pressoir, Geoff Morris.

## Abstract

Genomic mapping has yielded extensive catalogs of quantitative trait loci underlying agronomic traits, yet translating these discoveries into breeding gains remains inefficient. Here, we introduce Flywheel Genomics, a framework that integrates trait discovery directly within rapid cycling breeding populations. Using empirical data from a smallholder-oriented sorghum breeding program, we demonstrate that recurrent intermating and selection maintain genetic diversity, effective population size, and recombination while reducing confounding from plant height and maturity. Within this population, we resolve loci underlying simple adaptive and complex environmentally responsive traits and generate large segregating populations for mapping and near-isogenic lines for locus validation. We further demonstrate applicability in a public wheat breeding program, where known agronomic loci were readily detected. Simulations show that rapid cycling better preserves the population genetic properties required for Flywheel Genomics than conventional pure line development. By integrating discovery with improvement, Flywheel Genomics reframes breeding programs as engines of both crop improvement and genetic insight.

## INTRODUCTION

The effective translation of genetic discoveries into improved cultivars remains a central challenge in crop improvement. This challenge is particularly acute in low-input agricultural systems, where breeding gains must be achieved under resource and environmental constraints^1^. Although numerous quantitative trait loci (QTL) underlying key agronomic traits have been identified, their practical use in breeding remains limited^2^.

Trait dissection is typically conducted using synthetic populations such as biparental families, diversity association (AP), nested association mapping (NAM), and multi-parent (MAGIC) panels^3–7^. These panels are utilized across species, environments, and phenotypes to characterize genetic architecture and identify potentially economically important QTL^8–15^. These populations were designed to address specific challenges in genetic mapping; however, each carries inherent limitations. Biparental families are efficient to construct and can target specific traits, but capture only a limited portion of species-wide allelic diversity^16^. APs capture large portions of allelic diversity but exhibit strong population structure, which can confound association signals^17^. In addition, association mapping approaches tend to preferentially detect common, large-effect alleles, often missing rare variants^18^. NAM and MAGIC populations mitigate some population structure through recent recombination but require substantial time and resources to develop. Moreover, they may not segregate for traits relevant to a given target production environment^19–21^. In all these populations, variation in plant architecture phenotypes, such as flowering time and height, frequently complicates trait definition and introduces confounding effects^22,23^.

We argue that the principal limitation is not the mapping population itself, but the separation of genetic discovery from the breeding populations where these discoveries must ultimately be deployed. (Fig. 1a)^24,25^. Successful translation is contingent on several factors. Predictive molecular markers must remain accurate across genetic backgrounds^26^. Exotic germplasm may be required as donor material when alleles are absent, which may cause issues due to linkage drag, unfavorable background interactions, and context-dependent genetic effects that vary across environments and genetic backgrounds^27–29^. Moreover, revalidation in the target germplasm and environments is required, as genetic effects may vary across environments and genetic backgrounds^30,31^. As a result, QTL identified in static mapping panels often lose predictive value or exhibit reduced effects when deployed in breeding programs, limiting their utility.

**Fig. 1.**
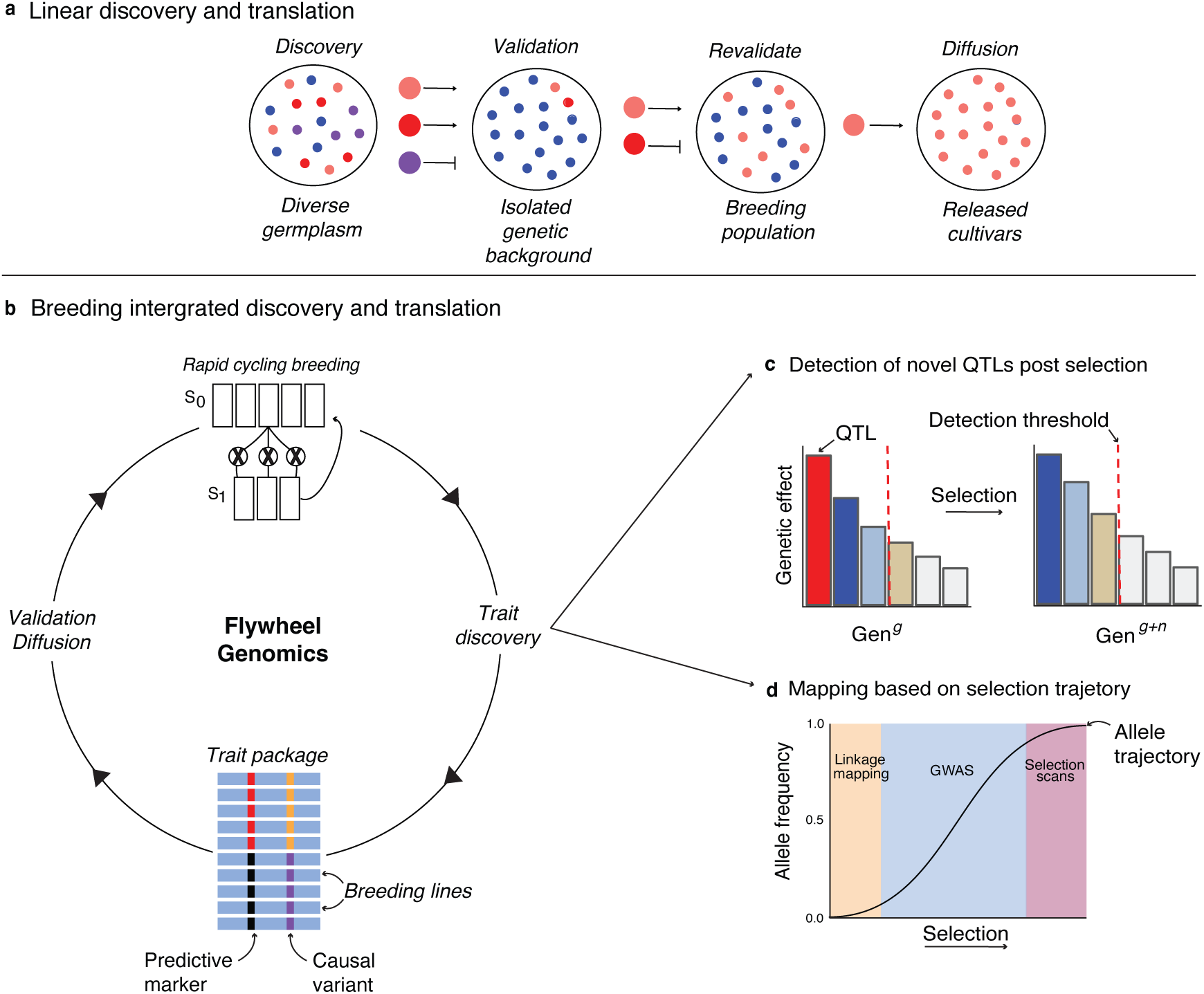
A shift from linear crop genetic translation to a dynamic breeding integrated Flywheel Genomics framework. **a**, Linear translation pipelines separate trait discovery, validation, and deployment, resulting in attrition of candidate loci. **b**, Flywheel Genomics integrates discovery, validation, and breeding within a rapid cycling breeding population. **c**, Directional selection reshapes genetic architecture across cycles, altering the set and effect sizes of detectable loci. **d**, The utility of mapping approaches depends on allele frequency: linkage mapping is most effective for rare alleles, GWAS for intermediate common alleles, and selection scans as alleles approach fixation.

The genetic architecture underlying complex traits is dynamic, changing over time as populations undergo selection and recombination. This is demonstrated by long-term selection experiments, such as the Illinois Long-Term Selection Experiment^32^. After more than 100 cycles of selection, maize populations continue to respond^33^, with loci shifting in effect sizes across cycles^34^, and genetic variance being continually reshaped rather than depleted^34^. These findings highlight that trait genetics are not static properties of a population but emerge through recombination and selection. Because most genetic studies are conducted in static populations, they inherently fail to capture these evolutionary dynamics. These limitations have motivated alternative strategies that more closely integrate trait genetics with breeding. One such approach, “map-as-you-go,” suggests QTL evaluation occurs directly within a breeding population^35^, which can reduce the gap between discovery and cultivar development by evaluating loci in the target genetic background and environment. Despite increasing use of modern breeding germplasm for GWAS^36–38^, no exploration of how breeding population design influences mapping power and resolution has been conducted.

We propose Flywheel Genomics (FG), a framework that leverages a highly intermated, rapid cycling breeding population in order to conduct trait genetic studies directly in line with breeding efforts. Realizing such a framework requires treating breeding populations as dynamic systems (Fig. 1b). Through cycles of recombination and selection, allele frequencies, linkage disequilibrium (LD), and trait architecture continuously shift^35,39,40^, and different QTL may be revealed and become detectable (Fig. 1c)^41^. Understanding this selection trajectory can inform the appropriate analytical approach to mapping^40^. Most importantly, the integration creates a self-reinforcing loop, in which each round of breeding generates new opportunities for discovery, and each discovery can accelerate the breeding (Fig. 1b).

Effective strategies for structuring breeding populations to support sustained genetic gain and trait discovery under strong directional selection remain largely unexplored. To explore this, a highly intermated sorghum (*Sorghum bicolor*) breeding population was developed. The Haitian Chibas sorghum breeding program uses rapid recurrent selection with a highly intermated crossing strategy to develop varieties for low-input smallholder agriculture under aphid pressure^42^. In 2015, a new sorghum aphid (*Melanaphis sorghi*) biotype emerged and devastated Haitian production, causing severe yield losses^43^. As a result, the gene pool reflects the effects of a severe bottleneck followed by population recovery, conditions expected to reduce genetic diversity, elevate LD, and lower heterozygosity^44^. Despite this, complementary mapping through selection scans and GWAS identified the resistance loci *RMES1* and *RMES2* (Fig. 3f; Fig. 4f)^45, 46^, demonstrating that adaptive variation can be detected within a breeding population post selection.

**Fig. 2.**
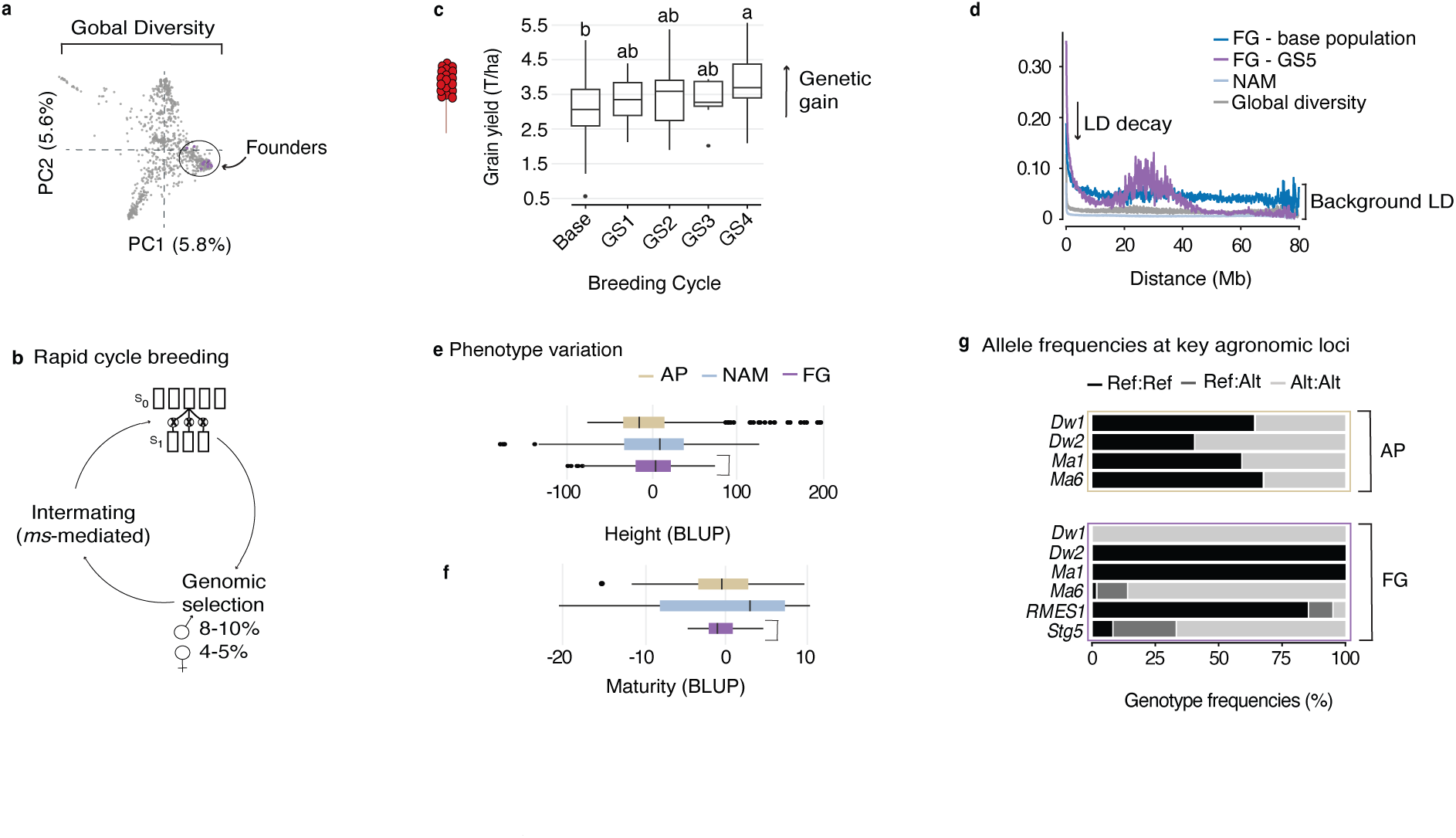
A highly intermated breeding population has favorable genetic properties for genetic gain and trait discovery. **a**, Principal component analysis showing Flywheel Genomics (FG) founders (purple) relative to global sorghum diversity (grey). **b**, The FG population (FGp) was established through rapid recurrent selection integrating genetic male sterility (ms3) and genomic selection (GS) across cycles. **c**, Grain yield across four GS cycles under high aphid pressure (Tukey’s HSD, P < 0.05). **d**, Linkage disequilibrium (LD) decay in the FG base population and GS5, compared to GDP and NAM panels. **e, f**, Distributions of plant height and maturity in the FG training population compared with NAM and an association panel (AP). f, Allele frequencies at key agronomic loci in AP compared to FGp .

**Fig. 3.**
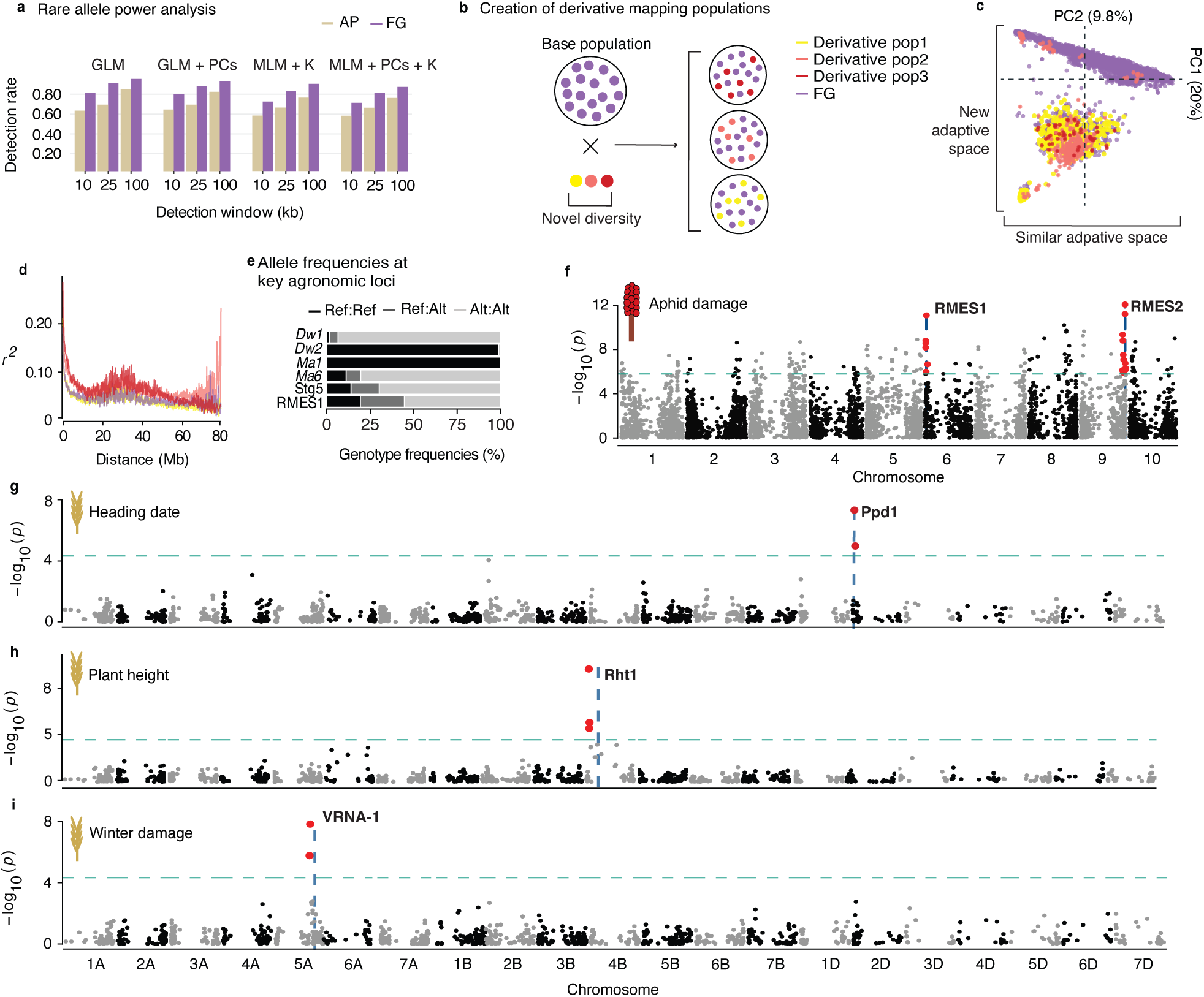
Intermating within an FGp generates powerful mapping populations from an adapted base population. **a**, Comparison of SNP–trait association detection rate for rare alleles (MAF between 0.05 and 0.06) between an AP and FGp. Detection was considered successful if a Bonferroni-significant SNP occurred within 10, 25, and 100 kb of the causal variant. Columns show the different models used with respect to the type of structure correction they include (PC = principal components; K = kinship). **b**, Derivative populations were generated by crossing the FG base population (purple) with diverse germplasm to introduce novel variation and target new adaptation (photoperiodic population; yellow; Guinea × IRAT 204 population; salmon; Haitian-Chibas landraces; red). **c,d**, PCA, and LD decay comparing derivative populations with base. **e**, Derivative population allele frequencies at key agronomic loci. **f, g, h, i**, GWAS Manhattan plots for sorghum and wheat traits. Red points indicate SNPs exceeding the Bonferroni genome-wide significance threshold (teal dashed line) within 10kb of a priori candidate loci.

**Fig. 4.**
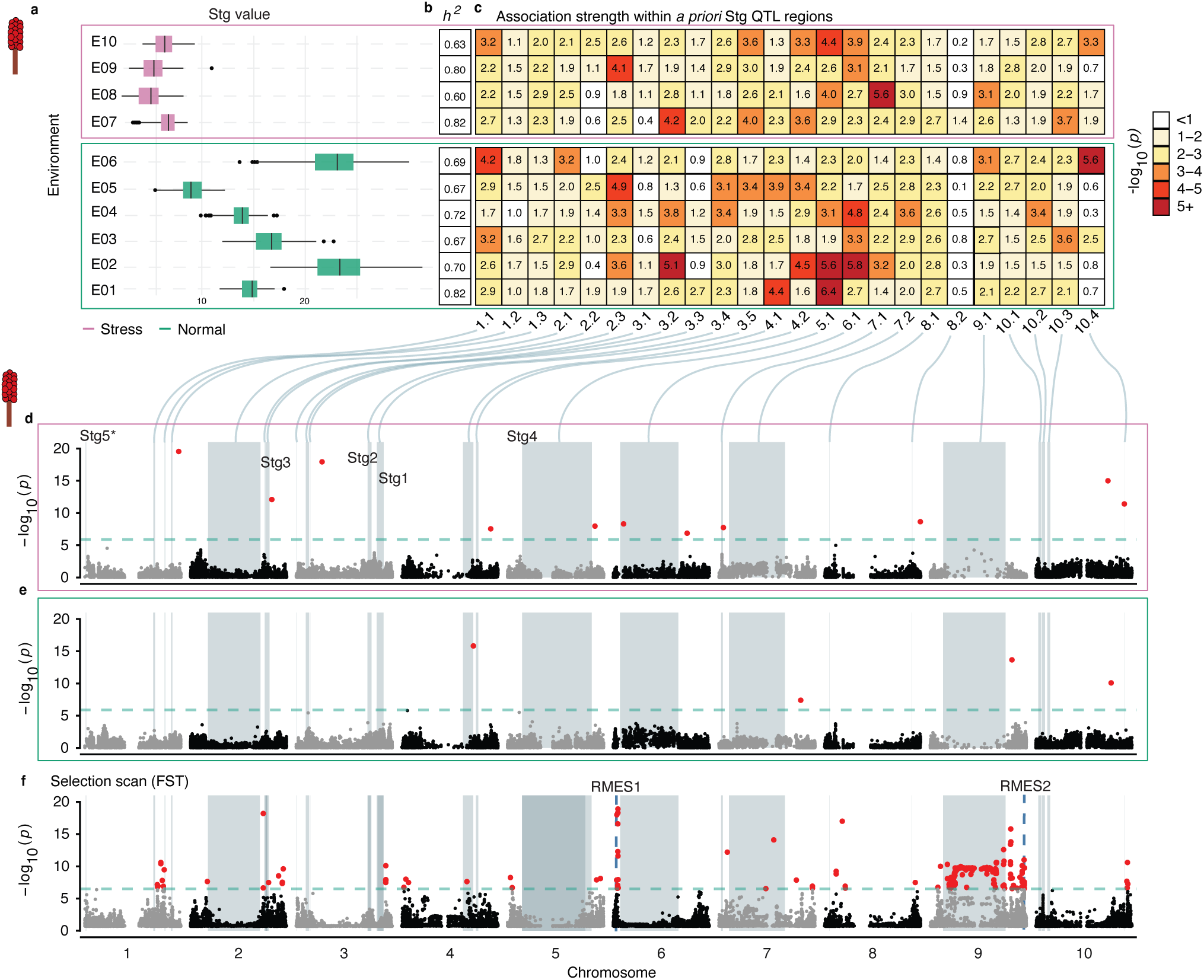
Multi-environmental GWAS in an FGp reveals the plastic nature of stay-green. **a**, Distribution of stay-green scores across ten environments, including non-stressed conditions and water- and salinity-stressed environments. **b**, Narrow-sense heritability estimates across environments. **c**, Heatmap of MLM GWAS -log10(p) values across environments of the peak-associated SNP within the boundaries of previously mapped stay-green QTL. **d,e**, BLINK GWAS Manhattan plots of mean stay-green value under stress and non-stress. **f**, Genome-wide FST between global diversity and the FGp. Dashed blue lines show the location of aphid resistant loci. Shaded regions indicate boundaries of previously mapped stay-green QTL. Red points indicate SNPs exceeding the Bonferroni genome-wide significance threshold of 0.05 (teal dashed line). No SNPs colocalized within 100 kb of Stg5.

Building on these insights, we establish the FG framework for designing breeding systems that support simultaneous genetic discovery and crop improvement. Using empirical data, we demonstrate how a rapid cycle breeding can achieve impressive genetic gain while exhibiting ideal mapping characteristics, as compared to static mapping resources. Through breeding simulation, we explore how alternative strategies influence genetic gain and mapping potential, thus providing the principles for designing FG populations.

## RESULTS

### Rapid cycling drives genetic gain while maintaining diversity

The primary objective of a breeding program is sustained genetic gain. We evaluated this using the inaugural sorghum Flywheel Genomics population (FGp), founded in 2013 from ten founders and designed for rapid recurrent intermating using genetic male sterility (*ms3*) (Fig. 2a,b)^42^.

Mean grain yield increased by an average of 75 kg ha⁻¹ per cycle across six environments, ranging from 30 to 148 kg ha⁻¹ (Supplementary Table 1). Assuming two selection cycles per year, this corresponds to an annual gain of approximately 150 kg ha⁻¹ yr⁻¹ (4.6% yr⁻¹), exceeding rates commonly reported for many breeding programs (Supplementary Table 2). Gains were particularly pronounced under severe sugarcane aphid pressure (Fig. 2c), demonstrating a strong adaptive response to selection.

Because breeding populations originate from a finite number of founders^49^ and experience repeated cycles of selection^50^, erosion of genetic diversity is a concern. Since the FGp has undergone strong directional selection, we tested whether diversity was eroding. From the base population to GS5, expected heterozygosity (*H*_e_) remained stable, and effective population size (*N*_e_) increased approximately threefold (Supplementary Table 3). Consistent with these patterns, the rate of LD decay increased and background LD decreased across cycles (Fig. 2d).

These results indicate that extensive intercrossing and rapid cycling can achieve high genetic gain while maintaining genetic diversity at non-target loci.

### A breeding population can exhibit ideal genetic mapping properties

Effective trait mapping requires a population with genetic diversity, high levels of effective recombination, and limited confounding variation^51^. One common source of confounding in crop studies is variation in height and maturity, which can influence many phenotypic measurements and obscure the detection of causal loci^22^. In breeding programs, these traits are considered attained because their reduced variation aligns with desired agronomic performance and farmer preferences^52,53^. Relative to established sorghum mapping panels, the FGp displayed a narrower range of height and maturity values and had less extreme values (Fig. 2e,f). Marker allele frequencies reflected this narrow distribution, as Kompetitive Allele-Specific PCR (KASP) genotyping indicated that the FGp was near fixation at major loci for height and maturity (Fig. 2f). Together, reduced phenotypic variance and fixation at major loci likely minimize confounding effects as compared to established mapping populations.

Alongside reduced phenotypic confounding, the extent of effective recombination influences mapping resolution by shaping linkage disequilibrium and the ability to fine-map causal loci^54^. For example, increased recombination from repeated intercrossing is expected to break down long-range haplotypes. To evaluate this property, we compared the rate of LD decay in the FGp to the sorghum NAM and a global diversity panel. Initial short-range LD decay occurred at comparable rates (Fig. 2g). However, the FGp maintained higher levels of intermediate and long-range LD, consistent with recent directional selection and selective sweeps^55^. However, this long-range LD was reduced in the GS5 generation compared to the base population (Fig. 2g), indicating that effective recombination is breaking down the extended haplotypes in a breeding selection context.

### Breeding populations can serve as platforms for genetic discovery

A FG breeding population can function as a platform for genetic mapping and variant validation. We compared locus detection rates in the FGp and an AP via simulation of low and high heritable (*h²*) single-locus traits with rare or common variants. Both detected common alleles at similar rates (Supplementary Fig. 1), even though the AP had a greater number of segregating SNPs. The FGp consistently had higher detection rates for rare variants (minor allele frequency range 0.05 - 0.06) for both high and low *h²* traits (Fig. 3a). The AP did detect common variants for low *h²* at a higher rate when using models that incorporated kinship (Supplementary Fig. 1c). This suggests that common alleles in the FGp may be more strongly confounded with relatedness, reducing their detectability when stringent correction is applied. Overall, these simulation results indicate that a FG approach can increase the detection of rare variants, which are often missed by GWAS in AP^18,56,57^.

For mapping of fixed loci, a key advantage of a FGp is the ability to rapidly generate derivative mapping populations directly from an active breeding program. Derivative populations can be tailored to specific breeding objectives while retaining the adaptive genetic background of the main FG population.^46^. Illustrating this, we crossed selected FGp lines with diverse germplasm (Fig. 3b). These derivative populations formed distinct genetic clusters while retaining evidence of shared ancestry with the main FGp, reflecting the recent admixture (Fig. 3c)^58^. Background LD and rate of decay remained similar to the base population (Fig. 3d), indicating that recombination patterns were preserved, in contrast to when structure is increased at population creation^59^. Marker genotyping further showed that these populations were segregating for previously fixed loci in the base population, while remaining largely fixed for major height, maturity, and stay-green (Stg) loci (Fig. 3e). Together, these features likely contributed to the remapping of the aphid resistance QTL, *RMES1* (Fig. 3f)^46^, which was fixed in the main FGp (Fig. 3g; Fig. 4f)^45^. Together, these results establish the genetic mapping potential of the FG framework.

### Breeding populations reveal environment dependent genetic architecture

We previously showed we can resolve loci for traits with relatively simple genetic architecture^46,60^. However, it remained unclear whether the FG approach could detect loci underlying more complex, environmentally responsive traits. To examine this, we utilized multi-environment phenotyping of the FG training population to perform GWAS for stay-green (Stg; delayed leaf senescence), under both drought-stress and non-stress conditions (Supplementary Table 4; Fig. 4a). Narrow-sense heritability ranged from 0.62 to 0.82, indicating differences across environments were due to underlying additive genetic components (Fig. 4b). GWAS was conducted separately for each environment as well as on the mean value across stress and non-stress conditions. Multiple significant loci colocalized with previously mapped Stg QTL (Fig. 4d,e). However, the significance of these loci varied across environments, with some detected only under drought stress and others under non-stress conditions (Fig. 4c). This pattern reflects environment-dependent genetic effects and may inform target environment breeding efforts^61^. Several previously mapped QTL did not have significant SNPs colocalizing but showed strong fixation signatures (Fig. 4g), highlighting the FG emphasis on utilizing different mapping tools based on the allele frequency for a trait under selection (Fig. 1d). Importantly, such QTL characterization was facilitated by the routine multi-environment testing intrinsic to breeding programs.

### Breeding populations can generate variant validation resources in relevant germplasm

While the FG approach can support locus detection, variant validation requires testing allelic effects independently of the genetic background. Although near-isogenic lines (NIL) are the standard approach for this^62,63^, their development is time and labor intensive. The heterogeneous inbred family (HIF) approach provides an alternative by leveraging residual heterozygosity within partially inbred lines^64^. Each HIF is mostly homozygous but segregates at the target region, allowing the extraction of NILs and comparison of alternative alleles in an otherwise similar genetic background. The FGp provides an opportunity to implement a HIF approach because it contains individuals with high levels of genome-wide fixation with retained heterozygosity at specific loci. Most loci were represented by NIL-like candidates, with 97% of markers having at least one candidate at a 0.7 threshold and 50% at 0.9 (Supplementary Table 5). These NIL-like candidates can be advanced through selfing to fix alternate alleles at target regions. These results provide empirical evidence to support an HIF approach using a FGp.

### Demonstrating Flywheel Genomics as a crop-agnostic approach

Building on the success in sorghum, we initiated a wheat rapid cycling population using commercially oriented elite germplasm. The population was derived from 26 breeding lines from the Kansas State University wheat breeding program and subjected to three rounds of random intermating before doubled haploid development. This created a partially recombined breeding population suitable for early-stage genetic analysis. GWAS for heading date, plant height, and winter damage identified significant associations that colocalized with known causative loci, including *Ppd1*, *Rht1*, and *VRNA-1* (Fig. 3g–i), demonstrating that FG provides a generalizable strategy for integrating genetic discovery into diverse crop improvement programs.

### Simulations reveal how rapid cycling enables simultaneous gain and discovery

To determine how breeding strategy influences the balance between genetic gain and trait discovery, we implemented forward-time simulations modeling recurrent selection under alternative crossing strategies. We compared pure line development (PL), which relies on repeated selfing with limited intercrossing, to rapid cycling, which implements extensive intercrossing each generation. In rapid cycling, females were generated via a recessive genetic male sterility (GMS) marker or randomly assigned, mimicking chemical sterilization (CS).

Selection targeted two traits: a highly polygenic trait representing yield and an oligogenic trait representing stress tolerance. Selection initially focused on yield (generations 1–10), after which a shift in selection pressure was imposed in which selected individuals required the favorable allele at a major-effect QTL for stress tolerance. In the remaining generations, a weighted selection index was implemented (80% yield, 20% stress tolerance; Fig. 5a; see Methods).

**Fig. 5.**
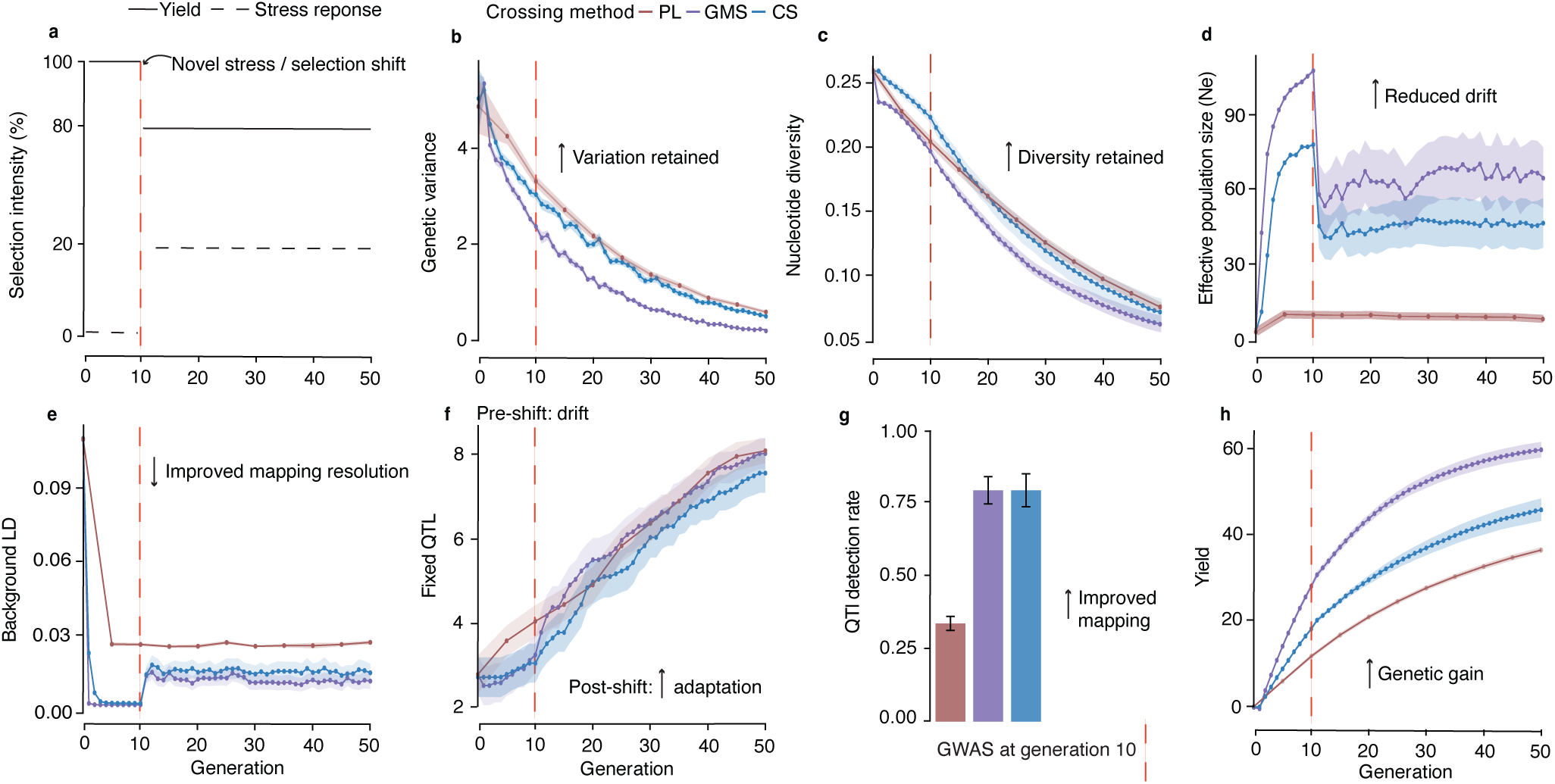
Rapid cycling strategies in an FGp sustain diversity and accelerate adaptation under shifting selection. **a**, A selection framework was imposed where initial selection was on a polygenic additive trait representing yield (solid line) for 10 generations. At generation 10 (red dashed line), selection was modified to consider a second oligogenic trait representing stress response (dashed line). **b**, Genetic variance for yield. **c**, Genome-wide nucleotide diversity. **d**, Effective population size. **e**, Background linkage disequilibrium (LD). **f**, Number of QTL fixed (MAF = 0) for the stress response trait. **g**, Average QTL detection rate for the stress response trait determined by GWAS at generation 10. Error bars represent standard deviation across replicates. **h**, Mean yield value.

We compared PL to GMS and CS to evaluate their suitability for FG. All three strategies experienced similar rates of decay in genetic variance and nucleotide diversity, although rapid cycling had slightly greater losses early on, possibly due to the increased number of selection cycles over a given time period (Fig. 5b-c). Rapid cycling maintained a larger *N*_e_ and lower background LD than PL development (Fig. 5d-e). Background LD remained low under rapid cycling despite increases in LD from selective sweeps, but not PL (Fig. 5e). Both *N*_e_ and LD decay in the rapid cycling mirror the observed trends in the FGp (Supplementary Table 3; Fig. 2d). These patterns are consistent with increased effective recombination and reduced drift from the higher numbers of crosses per generation compared to repeated selfing.

Breeding strategy also influenced the timing of QTL fixation and the ability to detect loci. PL development fixed alleles more frequently that either rapid cycling strategy before the shift in selection (Fig. 5f). In contrast, rapid cycling, particularly with GMS, accelerated fixation of favorable alleles after the shift in selection (Fig. 5f). Concerning mapping ability, rapid cycling substantially outperformed PL in QTL detection (Fig. 5g). These results indicate that extensive intercrossing not only delays premature fixation but also preserves the segregating variation necessary for effective QTL detection, while extensive selfing loses alleles to drift.

Finally, and most notably, rapid cycling had substantial genetic gain relative to PL development (Fig. 5h). The difference was more stark post selection shift. Overall, these simulation results suggest that while all strategies reduce diversity and increase desired traits, the rate and mechanism differ. This reflects the central advantage of the FG framework: QTL mapping can be conducted within a rapid cycling breeding population, accelerating the translation of genetic discoveries into crop improvement.

## DISCUSSION

Trait discovery via mapping approaches has previously been described as a “bandwagon” with limited relevance for plant breeding^65^. This argument may be largely due to the limitations and challenges in translating associations into improved cultivars, which arise from imperfect detection and the separation of discovery and deployment. Like a mechanical flywheel, which regulates the transfer of energy between a source and an engine, the FG framework links genetic discovery and crop improvement through a feedback process. In this context, mapping resolution is not fixed but emerges during population improvement. Our simulations provide support for how population development strategies differ in their outcomes, but they are necessarily simplified. Further exploration of parameter space, including selection intensity, crossing design, and population size, will be required to refine optimal strategies.

Our results support key features of the FG framework: that a population improvement approach to breeding can drive genetic gain and create a trait discovery platform. Importantly, even when large-effect loci are not detected, trait characterization within a breeding population generates actionable knowledge about genetic architecture, including polygenicity and genotype-by-environment interactions, which can guide selection strategies. Because these insights are derived within the target population, they are inherently translatable. Although demonstrated here primarily in sorghum, the framework is broadly applicable across crops, but adjustments will be needed depending on the mating system and constraints on pollination and crossing methods. The concept remains the same: a redefining of trait genetics as activity intrinsically embedded with breeding.

## MATERIAL AND METHODS

### Development and phenotypic evaluation of sorghum Flywheel Genomics population

The inaugural sorghum FGp (base population) was founded as S0-derived families generated by crossing superior S0 plants from the CIRAD-ms3 × ICSV 25280 background to Coloudo Nevado, 00-SB-FSDT-427, IS23563, WILEY, CIR-1/OG2-4G-1G-M-M, PCR-2>723C-1-M-1, Papesek, and Dekabes (IS 23572). Post aphid infestation, a training set was created by selecting a set of 296 representative individuals from the base population and selfing them to inbred status. The base population has since undergone five generations of GS by predicting genetic breeding values using the training population^66^. Genomic selection (GS) was applied with ∼8–10% of males and 5% of females being selected each cycle. Among selected individuals, approximately five random males were crossed to a random female using bulk pollination. The resulting offspring (designated as S_0_), along with select selfed males (S_1_), became the subsequent cycle.

Field trials were conducted using selected lines in the base population, and GS cycles one through four in four locations across Haiti from 2022 to 2024 (Supplementary Table 2). Genotypes were planted in randomized complete block designs with two blocks per year and location. In all environments, grain yield (tonnes/hectare) was measured for a total of 132 genotypes, including 85 from the base population, 13 from the first genomic selection cycle (GS1), 12 from the second GS cycle (GS2), 4 from the third GS cycle (GS3), and 8 from the fourth GS cycle (GS4). Grain yield was estimated as 85% of the panicle weight harvested at grain physiological maturity. Mean differences were compared using Tukey’s HSD implemented in the HSD.test() function of the agricolae R package. Days to anthesis and plant height were recorded at the Mirebalais location in 2024 (n = 732). Heading time was measured as the number of days from sowing to the date when approximately 50% of the plants in the plot had headed. Plant height was measured using a graduated ruler, from the base of the plant at the stem to the flag leaf. Best linear unbiased predictions (BLUP) were estimated with the model trait ∼ Block + (1|Genotype) using the R package lme4.

Stay-green (Stg) was assessed on the training population across 10 locations and year combinations (Supplementary Table 4). Fifteen days after physiological maturity, five plants were randomly selected per plot, and the plot-level Stg value was calculated as the total number of green leaves across the five sampled plants. Higher values reflect reduced senescence and improved retention of green leaf area. Per location genotype BLUPs were estimated with the model trait ∼ Block + (1|Genotype) using the R package lme4. GWAS was conducted on each environment independently using a mixed linear model that incorporated kinship^67^. The average of individual BLUPs across stressed and non-stress environments was used as the phenotype input for the GWAS algorithm BLINK^68^. All GWAS were run with GAPIT^69^. SNPs were colocalized with all Stg QTL reported in the sorghum atlas^70^.

### Genotype data

The global diversity panel (GDP) was previously characterized^60^. Briefly, a set of globally diverse germplasm (n = 1269), which included the sorghum FGp founders (n =10), was sequenced using genotype-by-sequencing (GBS) and aligned to the BTx623 sorghum reference genome v.3.1. The FGp training population data (n = 296) were obtained in the same GBS project.

Select lines from the base population (n = 210) and GS5 (n = 2072) were obtained from Diversity Arrays Technology genotyping (DArTAg). SNP data were quality-filtered for minimum call rates of 50% for both individuals and markers, reproducibility > 90%, and monomorphic sites, then mapped to chromosomal positions and converted from numerical genotype encoding to standard biallelic dosage format (0,1,2). The filtered data were reformatted to HapMap format using dartR^71^, converted to VCF using rTASSEL^72^, and subsequently imputed with Beagle v5.5^73^.

Derivative mapping populations were generated by crossing individuals from the FG base population with diverse sorghum germplasm. The previously published DArTseq genotype dataset generated for the FG base population (n = 3863) and derivative populations (photoperiodic population n = 399; Guinea × IRAT 204 population n = 319; Haitian-Chibas landraces n = 48) has been described^46^. Genotype calling, marker filtering, genomic alignment, and imputation were performed as described in the original study. The final SNP dataset was used for PCA, LD estimation, and GWAS presented here.

### KASP marker genotyping of known adaptive loci

Randomly selected samples from the base population (n = 367) and derivative populations (n = 368), as well as the accessions from the Sorghum association panel^74^, were sent to Intertek for Kompetitive allele-specific PCR (KASP) genotyping using allele-specific primers for target SNPs. KASP markers for *RMES1* were previously generated and characterized as predicting aphid resistance^45^. Markers associated with *Dw1* and *Dw2* were previously described^75^. KASP markers were designed for tagging maturity genes *Ma1*^76^ (*ma1-milo* allele, Sbv3.1_06_40306243 G/-) and *Ma6*^76^ (ma6-1 allele, Sbv3.1_06_00697740 -/GTGCA). SNPs with >5% MAF in the 100-bp flanking sequence of the desired allele were identified using the sorghum pangenome ^77^, and KASP primers were designed to avoid these polymorphic regions to minimize the risk of amplification failure from sequence variation at the primer binding site. STG5 markers that targeted the putative causal variant SNP (Sbv3.1_01_01144852 C/T) in the CYP79A1 gene and a SNP in LD (Sbv3.1_01_01287238 C/T) with the CYP79A1 gene that was private to Staygreen donor BTx642 were designed. Genotypic discrimination of the markers was validated using sorghum lines with the favored allele and lines with alternate alleles. For all KASP markers, genotypes were called and returned via Intertek’s standard software.

### Published diversity panel data

Two sorghum mapping panels were compared to the FGp. The NAM population was previously developed by crossing 11 diverse male founder accessions with the recurrent female parent ‘Grassl’. The population is also referred to as the carbon-partitioning NAM, as these 11 male founders were selected from the larger sorghum Bioenergy Association Panel based on their broad diversity in carbon-partitioning and bioenergy-related traits^78^. In the summer of 2020, phenotypic data for traits were collected at multiple sites in Pendleton, South Carolina^79^. The Sorghum Association Panel (AP) was developed to provide a genetically and phenotypically diverse collection of sorghum accessions suitable for association mapping^80^. Phenotype data used in this study were from a prior study in which field evaluations were conducted in Lincoln, Nebraska during the years 2010 through 2019^81^. Data from additional locations were also available; however, they all showed similar distributions and therefore only Nebraska was selected to be representative in this study. For both the NAM and AP, BLUPs were estimated for days to anthesis and plant height using linear mixed-effect models in the lme4 package in R. Each trait was analysed with a model that included replication as a fixed effect and genotype as a random effect, in the form trait ∼ Rep + (1|TAXA). For both populations, the preprocessed and filtered GBS data from the prior studies were incorporated and used in all analyses reported here.

### Population genetic analyses

LD decay was estimated using the LD.decay function from the sommer R package^82^, which computes pairwise correlation (*r^2^*) values between markers and their physical distances across all chromosomes. Markers were thinned to every fifth SNP to reduce computational burden, and *r^2^* values were binned into 10 kb distance intervals and fitted with a loess curve to visualize decay patterns across panels. Principal component (PC) analysis was conducted using the prcomp function in R. Genotypes were centered and scaled prior to computing. To identify the signatures of selection post hard sweeps from aphid pressure, a previously conducted fixation index scan (FST) between the FG training population and GDP was replotted for this study^60^.

### Genomic inbreeding coefficient estimation

To compare diversity and identify near-isogenic line (NILs) candidates, we used the base population (n = 210) and GS5 (n = 2072) DArTAg data. Markers with excessive missing data were filtered using a threshold of 50% maximum missingness per marker. Missing genotypic values were imputed using the mean allele frequency method implemented in the A.mat() function of the rrBLUP package. Individual genomic inbreeding coefficients (F) were estimated using marker-based measures derived from observed and expected heterozygosity. For each SNP, allele frequency (*p*) was calculated as half of the mean allele dosage across individuals. Expected heterozygosity (Hₑ) was computed as: *H*e = 2*p*(1 − *p*), and observed heterozygosity (H_0_) for each individual was calculated as the proportion of heterozygous loci across all markers. The individual inbreeding coefficient (F) was then estimated as: 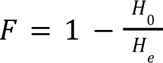.

### Identification of NILs across inbreeding thresholds

To characterize the distribution of residual heterozygosity across the genome, individuals from GS5 (n = 1,935) were filtered using a series of increasing inbreeding thresholds (F > 0.0 to F > 0.9, in increments of 0.1). For each threshold and population, only individuals exceeding the specified F value were retained. Within each filtered subset, SNP markers were screened to retain only polymorphic loci, defined as markers exhibiting more than one allelic state among the retained individuals. For each polymorphic marker, the number of heterozygous individuals was counted. This count was interpreted as the number of NIL candidates at a given locus, as each heterozygous individual could be selfed to generate progeny segregating for alternative homozygous alleles and used to develop a NIL pair.

### Genome-wide association studies QTL detection rate analysis

GWAS detection rates were compared between the FG training population (n =296; 6,841 SNPs) and the Sorghum AP (n = 307; 20,977 SNPs) using simulated single-locus phenotypes. To standardize marker density and reduce computational burden, analyses were restricted to chromosome one. To simulate monogenic traits, a single additive quantitative trait nucleotide (QTN) was randomly selected from the available SNP set and assigned an effect size of 1.0. Simulated phenotypes were generated using the simplePHENOTYPES^83^. To simulate a realistic GWAS setting in which the causal variant is not directly observed, the simulated QTN was then removed from the genotype dataset prior to association analysis, requiring detection via LD with neighboring markers.

To compare rare and common SNP detection in low and moderate heritable traits, four simulation scenarios were evaluated. (i) h² = 0.5 without minor allele frequency (MAF) constraint; (ii) h² = 0.5 with the causal QTN constrained to a MAF 0.05–0.06; (iii) h² = 0.3 without MAF constraint; and (iv) h² = 0.3 with the causal QTN constrained to a MAF 0.05–0.06. In unconstrained scenarios, QTNs were selected from any polymorphic SNP. In constrained scenarios, QTNs were selected only from SNPs within the specified MAF range. For each population and scenario, 100 independent replicates were simulated. Four GWAS models were implemented: (i) GLM (no correction for population structure), (ii) GLM including the first three principal components (PCs) as fixed effects, (iii) MLM including a random effect modeled using a precomputed kinship matrix (K), and (iv) MLM including both PCs (fixed effects) and kinship (random effect). All models were run using GAPIT^69^. Bonferroni correction was applied to control for multiple testing (α = 0.05 / number of SNPs tested). A replicate was considered successfully detected if at least one SNP exceeded the Bonferroni significance threshold and fell within a defined window (10, 25, or 100 kb) upstream or downstream of the simulated QTN position. Detection rate was calculated as the proportion of replicates (/100) meeting this criterion.

### Creation of Wheat Genomic Assisted Recurrent Selection (ProGARS) population

The wheat ProGARS population was developed from an initial set of 26 founder parents. The founder parents included germplasm of 2 high-protein spring wheats, 4 *Aegilops tauschii* derivatives, 4 *Triticum dicoccoides* derivatives, 7 experimental lines, and 9 released varieties. The parents were selected for their yield, quality, and/or protein deviation characteristics. All 26 founder parents were crossed to Bob Dole and Kanmark, both of which were previously backcrossed to incorporate the wheat ms3 male sterility gene^84^. The resulting F1 progeny were randomly intermated. After intermating, the female plants were tagged and harvested. The harvested seed was planted and randomly intermated again until a total of three intermating rounds were done. Following the third intermating round, the seeds were sent to Heartland Plant Innovations (HPI). At HPI, the males were selected and developed as double haploid lines. The resulting double haploid lines are the current ProGARS population.

### Wheat ProGARS Phenotyping

ProGARS genotypes were evaluated in Russell and Riley counties, Kansas, during the 2025 field season for plant height and heading date. The Russell environment included approximately 50% of the ProGARS genotypes evaluated due to limited seed availability and was excluded from the analysis due to the presence of Wheat Streak Mosaic Virus. Additionally, the Riley 2026 environment was evaluated for winter damage. Both environments were evaluated using a single unreplicated diagonal arrangement experimental design. Checks included Doublestop CL+, KS Mako, Bob Dole, and WB4699, systematically arranged in a diagonal distribution across the field. Each plot consisted of a 3-row plot with a total plot width of 0.60 m and length of 1.95 m. There was 1 m of spacing between ranges and 0.30 m of spacing between plots within the same range. The plot plant height was evaluated by measuring the average height for each plot. Plant height measurements were taken with a 240 cm long PVC pipe with measurement intervals every 2.5 cm starting from 50 cm to 240 cm. The heading date for each plot was recorded as the date on which ≥ 50 % of each plot had the wheat head emerged from the leaf sheath of the flag leaf. Heading dates were transformed into Julian dates prior to the GWAS analysis. Winter damage for each plot was evaluated using a 1-9 visual scale. For the visual scale, a 1 would be a plot with no visible damage, while a 9 would be complete winter damage to the wheat plants in the plot.

### Wheat ProGARS GWAS methodology

The genotyping platform was a set of 3996 SNP markers from the Agriseq Thermo Fisher Targeted GBS for wheat. A set of ProGARS experimental genotypes (n =342) was used. Twenty-six markers with unspecified chromosome and/or physical position were removed.

Eleven double haploid lines that were genotyped but not evaluated in the 2025 field season were also discarded prior to filtering. The set of 3970 markers and 331 genotypes was then filtered for MAF (0.05), call rate (0.95), and marker and genotype heterozygosity (0.1). Missing values were <2% of total marker data and were replaced with mean imputation. These filtering steps and imputation procedures were done using the snpReady version 0.9.7 R package^85^. The remaining markers were filtered for LD by the correlation method with a threshold of 0.2 and a sliding window of 250 kbp using SNPRelate^86^. After filtering, a total set of 1230 SNP markers and 299 genotypes was used for the GWAS analyses for plant height, heading date, and winter damage. Once the phenotype and genotype data were ready to be analyzed, the GWAS analyses were conducted using the MLM and multiple loci mixed model (MLMM) models in GAPIT^69^.

### Breeding simulations

To evaluate alternative strategies for population improvement, we simulated three breeding strategies: rapid cycling using either genetic male sterility (GMS) or chemical sterilization (CS), and pure line development (PL) where individuals are selfed to fixation prior to reuse. Across strategies, the best efforts were made to keep the number of crosses and the total population size equivalent (Supplementary Fig. 2). This allows us to compare population development strategies using consistent resource constraints.

### Breeding simulations: Creation of founder population

A genome was simulated with 10 chromosomes (total size = 10^8^ bp), a mutation rate of 2.5x10^-8^, and *N_e_* of 25. Using these settings, a pool of 300 diverse, heterogeneous individuals was created using forward-based coalescent simulation^87^. From this initial diverse pool, a set of 10 was randomly selected as population founders.

Population founding differed among the three breeding strategies. For GMS, an initial crossing stage was conducted where 9 of the selected founders were crossed with the remaining founder, which was heterogeneous for the *ms* allele. This facilitated the segregation of *ms* in the founding population. From each cross, 50 offspring were generated. From these offspring, 625 random crosses were simulated, and four offspring per cross were generated to create an initial population of 2500 plants. For the CS approach, all founders were randomly crossed for a total of 625, with four offspring per cross, to create the initial population of 2500 plants. For PL development, the initial population was developed by first randomly intermating the founders for a total of 625 crosses with four offspring from each cross for a total population of 2500. A single plant was selected from the 25 highest yielding F1 families and subsequently selfed to F5.

### Breeding simulations: Recurrent selection population development

In the rapid cycling strategies, 125 females and 250 males were selected each cycle based on their yield genetic value. Female sterility was imposed either through a recessive genetic *ms* locus (GMS) or by random assignment, mimicking CS. After selection, five males were randomly selected and crossed to a single female. This was done five times for each selected female. Four offspring were then randomly selected from each cross to be the next generation. Thus, each generation consisted of 625 crosses with 2,500 progeny.

PL followed a five-year breeding cycle. In year 1, 125 biparental crosses were generated by crossing each of the 25 parents with five random parents from the same pool. In year 2, 20 plants from each family were evaluated, and the two best individuals per family were advanced by selfing. In year 3, 10 plants per family were evaluated, and one plant from each of the top 125 families was selfed. In year 4, 20 plants per family were evaluated, and one plant from each of the top 50 families was selfed. In year 5, 50 plants per family were evaluated, and one plant from each of the top 25 families was selected. These 25 F5 lines were then intermated to initiate the year 1 breeding cycle. All three breeding strategies were run for 50 years and replicated 30 times.

### Breeding simulations: Trait simulation, selection index and population monitoring

Using the functions addTraitADEG and addTraitAD in the AlphaSimR package, we simulated two quantitative traits with different genetic architectures. Trait one was polygenic and represented yield, with 100 QTL per chromosome (1000 total), heritability 0.5, with normally distributed additive (mean = 0, variance = 5), dominance (mean degree of dominance set to 0.3), additive-by-additive epistasis (relative epistatic variance set to 0.3) effects, and genotype-by-environment interactions (variance = 2). Trait two was oligogenic with one QTL per chromosome (10 total), heritability 0.5, with additive effects (mean = 0, variance = 1) and dominance (mean degree of dominance = 0.5). Across all strategies, generations 1–10 were selected solely on Trait 1. To mimic an emergence of a novel selection pressure, for generations 11 onward, only individuals homozygous for the positive alleles for the largest effect QTL for Trait two were considered for advancement. For generations 11 onward, a weighted selection index (80% Trait one, 20% Trait two) was also employed (Fig. 5a).

### Breeding simulations: Simulation Metrics Collection

For all breeding strategies, baseline metrics were recorded once in the founding population (Generation 0) prior to the initiation of selection. In PL, metrics were subsequently collected at the beginning of each recurrent cycle following the formation of the F1 population. In rapid cycling, metrics were recorded once per cycle on the selected individuals. Population metrics, including phenotypic means, genetic variances, allele frequencies, background LD, and effective population size (Ne), were recorded. Genetic variance was estimated as the variance in individual breeding values using the AlphaSimR function varG(). Population response to selection was assessed using the mean phenotypic value for trait one using the function meanG(). Effective population size (N_e_) was estimated using a LD–based method^88^. N_e_ was estimated using the formula:

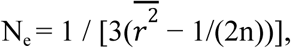

Where 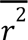 is the average pairwise LD between unlinked markers (located on different chromosomes), and n is the number of sampled individuals. *N_e_* was calculated using a random subset of 30% of total markers, excluding those monomorphic. The same estimate of 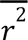 was also used as the measure of background LD. Nucleotide diversity was calculated per site as 2p(1−p), where p is the allele frequency, and then averaged across all segregating loci. Allelic frequency at causal loci for Trait two was quantified by calculating the minor allele frequency (MAF). Loss of variation at these loci was tracked by recording the number of QTL that reached fixation (MAF = 0).

## AUTHOR CONTRIBUTIONS

B.R., G.M., and G.P. conceived the Flywheel Genomics framework. G.P. and J.R.G. developed and managed the sorghum breeding population and collected field data. A.F. and E.M. developed the wheat rapid cycling population and generated phenotypic data. B.R., E.O., and J.R.C. performed genomic analyses and simulations. SRM and TF developed the KASP markers. B.R. interpreted the data, generated visualizations, and wrote the manuscript with input from all authors. All authors reviewed and approved the final manuscript.

## DATA AVAILABILITY

All data needed to evaluate the conclusions in the paper are present in the paper as appropriate citations to previously published data or included in the Supplementary Materials.

## CODE AVAILABILITY

R scripts used to perform the breeding and GWAS simulation studies are available at github.com/ebenogoe/flywheel-genomics-simulation.

## SUPPLEMENTARY FIGURES AND TABLES

**Supplementary Fig. 1.**
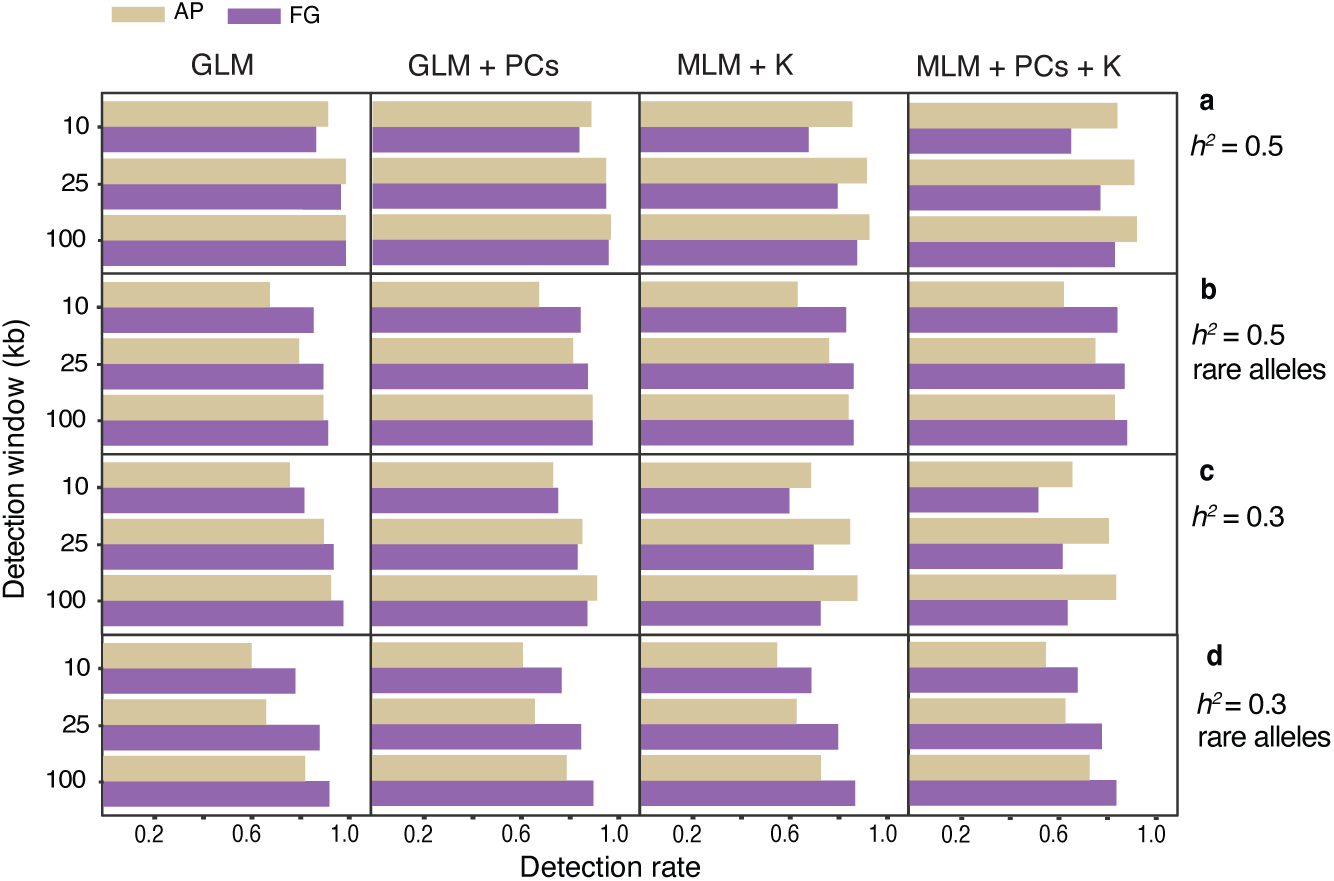
Variant detection rates comparing the FG population with an association panel (AP) using simulated genetic architectures. Detection rate of SNP–trait associations in the FGp and an AP across simulated traits with a single causal variant. Detection was considered successful if a Bonferroni-significant SNP occurred within 10, 25, and 100 kb of the causal variant. Columns show models (GLM, GLM + PCs, MLM + K, MLM + PCs + K). Rows correspond to simulated genetic architectures using different narrow-sense heritabilities: a, *h²* = 0.5; b, *h²* = 0.5 with rare alleles (MAF 0.05–0.06); c, *h²* = 0.3; d, *h²* = 0.3 with rare alleles. Detection rate is reported as the number of times /100 independent runs, a SNP was detected.

**Supplementary Fig. 2.**
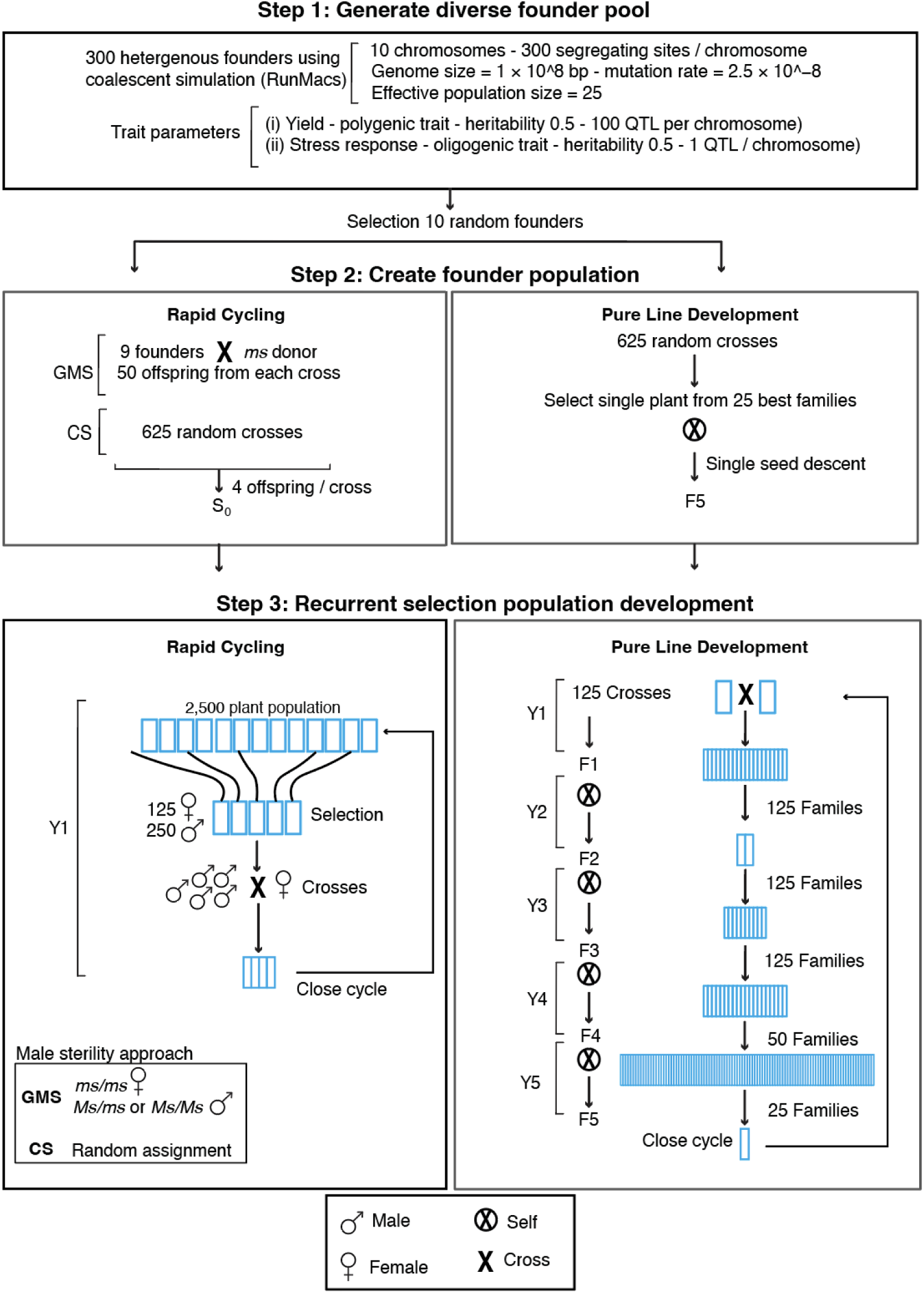
Simulated Population Development Descriptions. Three breeding strategies were simulated: rapid cycling using genetic male sterility (GMS), rapid cycling using chemical sterilization (CS), and pure line development (PL). Each cycle began with 2,500 individuals derived from 625 crosses. Rapid-cycling strategies recycled selected progeny as parents every generation, whereas PL advanced families to the F5 generation before selecting 25 elite lines as parents for the next cycle. Founder populations were simulated in RunMacs. Traits included a polygenic yield trait and an oligogenic stress-tolerance trait. See material and methods “Breeding simulation” for full details.

**Supplementary Table 1.** Realized genetic gain for grain yield in FGp across three Haitian environments. Estimates are shown per selection cycle and per year, assuming two breeding cycles annually. Locations and year combinations are noted for environmental stress.

| Site | Year | Coordinates | Irrigation | Stress | $\Delta G$ kg<br>cycle<sup>-1</sup> | $\Delta G$ kg<br>year<sup>-1</sup> | $\Delta G$ %<br>cycle<sup>-1</sup> | $\Delta G$ %<br>year<sup>-1</sup> |
| --- | --- | --- | --- | --- | --- | --- | --- | --- |
| Cabaret | 2024 | 18°43'33.28"N,<br>72°25'3.241"W | Irrigated | - | 30.35 | 60.7 | 0.73 | 1.47 |
| Saint<br>Michel | 2023 | 19°25'10.52"N<br>72°20'33.67"W | Irrigated | - | 30.77 | 61.55 | 0.6 | 1.2 |
| Saint<br>Michel | 2023 | 19°25'10.52"N<br>72°20'33.67"W | Rainfed | Drought | 69.67 | 139.33 | 1.53 | 3.05 |
| Mirebalais | 2024 | 18°53'17.04" N<br>72°05'04.79" W | Irrigated | - | 63.27 | 126.54 | 2.73 | 5.47 |
| Saint<br>Raphael | 2022 | 19°22'21.09"N<br>72°10'32.05"W | Irrigated | - | 110.34 | 220.68 | 3.46 | 6.91 |
| Cabaret | 2022 | 18°43'33.28"N,<br>72°25'3.241"W | Irrigated | High aphid<br>pressure | 147.73 | 295.46 | 4.73 | 9.46 |
| **Mean** |  |  |  |  | **75.35** | **150.71** | **2.3** | **4.59** |

**Supplementary Table 2.**
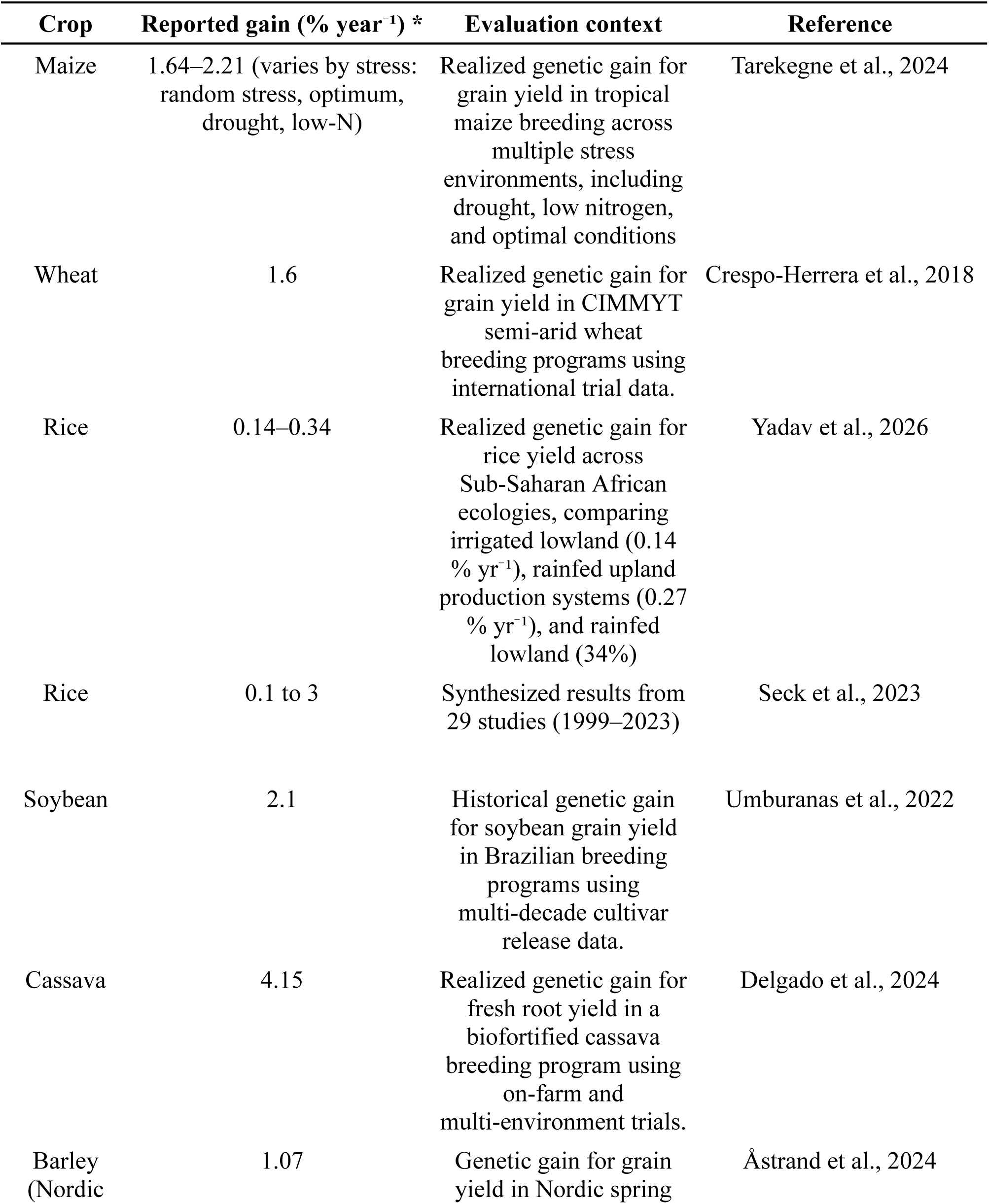
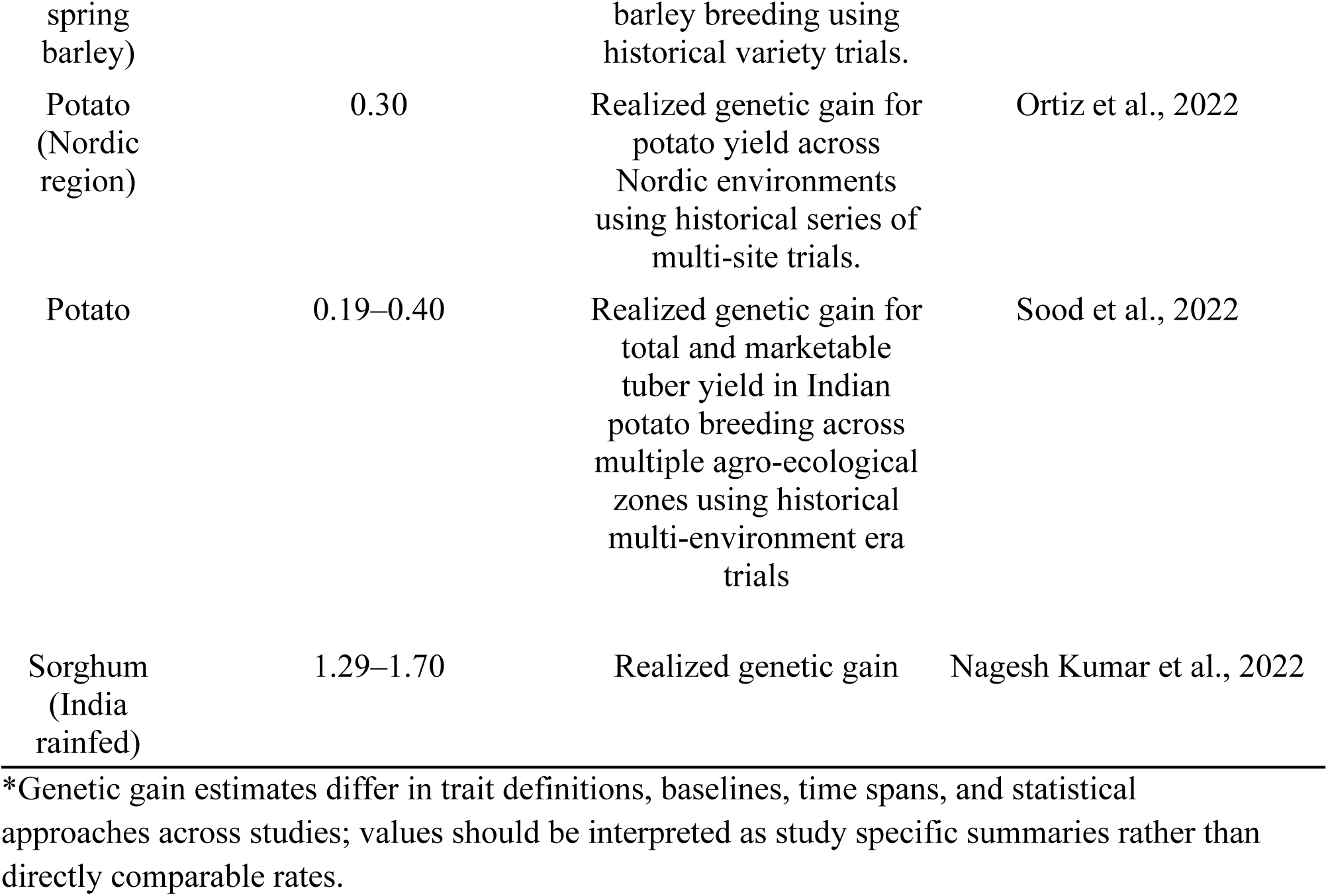
Summary of published estimates of genetic gain for grain or yield-related traits across major commercial and subsistence crops, expressed as percent improvement per year (% yr⁻¹).

**Supplementary Table 3.**
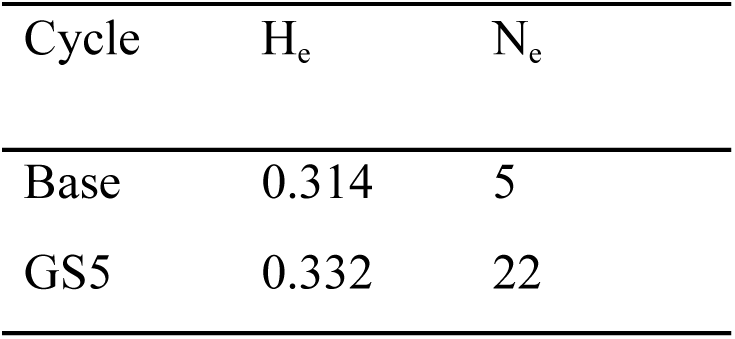
Diversity metric of the base and GS5 generations. Comparison of expected heterozygosity (*Hₑ*) and effective population size (*Nₑ*) between the base population (n = 98) and the fifth cycle of genomic selection (GS5; n = 172).

| Cycle | $H_e$ | $N_e$ |
| --- | --- | --- |
| Base | 0.314 | 5 |
| GS5 | 0.332 | 22 |

**Supplementary Table 4.** Location and year combinations of where the FG training population was evaluated for stay-green.

| **Environment** | **Site** | **Planting Date** | **Coordinates** | **Irrigation** | **Stress** |
| --- | --- | --- | --- | --- | --- |
| E01 | Cabaret | March/25/2017 | 18°43'33.28"N<br>72°25'3.241"W | Irrigated | - |
| E02 | Cabaret | May/8/2017 | 18°43'33.28"N<br>72°25'3.241"W | Irrigated | - |
| E03 | Cabaret | July/12/2017 | 18°43'33.28"N<br>72°25'3.241"W | Irrigated | - |
| E04 | Route9 | Sept/7/2017 | 18°38'54.56"N<br>72°17'59.43"W | Irrigated | - |
| E05 | Route9 | Nov/9/2017 | 18°38'54.56"N<br>72°17'59.43"W | Irrigated | - |
| E06 | Route9 | Oct/23/2018 | 18°38'54.56"N<br>72°17'59.43"W | Irrigated | - |
| E07 | Route9 | Nov/9/2017 | 18°38'54.56"N<br>72°17'59.43"W | Rainfed | Drought |
| E08 | Route9 | Feb/3/2018 | 18°38'54.56"N<br>72°17'59.43"W | Rainfed | Salinity/Drought |
| E09 | Route9 | Feb/16/2018 | 18°38'54.56"N<br>72°17'59.43"W | Rainfed | Salinity/Drought |
| E10 | Route9 | April/25/2018 | 18°38'54.56"N<br>72°17'59.43"W | Irrigated | Salinity |

**Supplementary Table 5.** Availability of candidates for near-isogenic line (NIL) locus validation per marker in the GS5 population. Mean, median, and quartile (Q) distribution of NIL per SNP marker in GS5 across genomic inbreeding thresholds (F), with the number of individuals retained at each threshold (n) indicated.

| F | n | Mean | Q1 | Median | Q3 | Markers represented by ≥1 NIL (%) |
| --- | --- | --- | --- | --- | --- | --- |
| 0.7 | 806 | 44.08 | 10 | 22 | 52 | 97% |
| 0.8 | 529 | 21.11 | 1.75 | 7 | 20 | 82% |
| 0.9 | 258 | 7.19 | 0 | 0 | 3 | 48% |

